# fingertipsR: an R package for accessing population health information in England

**DOI:** 10.1101/189167

**Authors:** Sebastian Fox, Julian Flowers, Simon Thelwall, Daniel Flint, Doris Hain

## Abstract

Fingertips is a major public repository of population and public health indicators for England, built and maintained by Public Health England (PHE). The indicators are arranged in thematic or topical profiles covering a wide range of health issues including:

- broad Health Profiles
- specific topics such as liver disease and end of life
- risk factors including alcohol, smoking, physical activity
- population healthcare health services data for general practices, cancer, mental health
- health protection data on general health protection, TB, antimicrobial resistance
- lifecourse profiles for younger and older people
- mortality and morbidity.

Fingertips makes data available for more than 1,500 indicators spread across more than 70 profiles. The data can be accessed from https://fingertips.phe.org.uk where the data are visualised in variety of ways including heatmaps, choropleth maps, line charts for trends, “spine” charts (graphs which compare multiple indicators for a single geographic area), scatter plots and so on. Data can be obtained as downloads or figures which can be exported or cut and paste into reports and slides.

A recent addition to the Fingertips platform was an Automated Programming Interface (API) to enable developers to re-use the data. To facilitate access to Fingertips data we have designed an *R* package - `fingertipsR` - enabling rapid and easy access to the data by analysts and data scientists. The package is available from the Comprehensive R Archive Network (CRAN).

This paper describes the `fingertipsR` package which provides tools accessing a wide range of public health data for England from the Fingertips website using its API.

## Introduction

Fingertips is a major public repository of population and public health indicators for England, built and maintained by Public Health England (PHE). Data are organised thematically, grouping related indicators together. The web interface provides an interactive platform for data visualisations allowing users to examine trends over time, geographical distribution and make comparisons between health providers. The web interface also provides a means to download the data behind the visualisations for re-use. However, accessing the data in this manner limits the user to a single group of indicators for a specified geography per download. To enable programmatic access to the data PHE developed an application programming interface (API). However, use of an API is highly technical and not always suited to the public health researcher.

R is a free, open source software for statistical analysis. (R Core Team 2017) It doubles as both a programming language and analytical environment for performing statistical analyses. The programming language facilitates expansion of the software through additional user-written ‘packages’ which are then stored in an online repository. Such packages bundle together analytic commands which share a common purpose. The ease with which R can be expanded through its packaging system has led to exponential growth in the software, creating a very broad ecosystem of statistical techniques. (J. Fox 2009)

The `fingertipsR` package extends R by providing an easy-to-use set of functions to query the Fingertips API, allowing direct import of data to R. (S. Fox 2017)

## The fingertipsR package

### The structure of data in fingertips

Public health data gathered and synthesised by PHE are stored on Fingertips in nested thematic groups. Profiles group together broad themes of data such as antimicrobial resistance or diabetes. (Johnson et al. 2016) These profiles may consist of multiple domains - such as prevalence or targets. Individual indicators then provide actual values for different measures within the domains for example prevalence among those ≥ 65 years of age. In addition, indicators can vary by different area types within two broad geographic themes, administrative and health. Health geographies include commissioners of health care services for local areas (known as Clinical Commissioning Groups or CCGs), hospital groups (acute trusts) and general practices. Administrative geographies include upper- and lower-tier local authorities and government regions among others. These geographies themselves fall within nested geographies based on whether they are in the administrative or health geography hierarchy.

All indicators have a fundamentally identical structure. Columns provide: a code that uniquely identifies a geography, three time variables that specify the year, quarter and month, counts and denominators (data are provided where appropriate) and a value column gives the actual value - such as a prevalence or rate - to be plotted in the various representations on the website. Rows provide an observation for the indicator at a specified time period, age group and sex for a given geography.

### Accessing the fingertips API

The API aims to provide the same data through a collection of RESTful web services as can be visualised on the Fingertips web site. The API was created following open data principles so that the data on the Fingertips site could be available for anyone to access, use or share. (Ahmed 2012) The web-based API enables the data to be accessed from any location over the internet using the researcher’s programming language of choice without any knowledge required of the system that provides that data. The data are transferred in JavaScript Object Notation (JSON), a lightweight data format that is intended to be easy for both humans to read and machines to generate and parse.

### Functions provided by the fingertipsR package

The functions of the fingertipsR package facilitate exploration of the fingertips data in a way that reflects the structure of the data. A public health researcher may start by examining which profiles and domains are available:

~~~
library(fingertipsR)
*# for common data manipulation functions*
library(dplyr, warn.conflicts = FALSE, quietly = TRUE)

## Warning: package ’dplyr’ was built under R version 3.3.3
ftips_profiles <- profiles(ProfileID = NULL, ProfileName = NULL) head(ftips_profiles)

## ProfileID     ProfileName     DomainID
##1     8 Adult Social Care 1000101
##2     8 Adult Social Care 1000102
##3     8 Adult Social Care 1000103
##4     8 Adult Social Care 1000104
##5     8 Adult Social Care 1000105
##6     8 Adult Social Care 1938132733
##                             DomainName
## 1     Enhancing quality of life for people
## 2 Delaying and reducing the need for care and support
## 3 Ensuring a positive experience of care and support
## 4                     Safeguarding vulnerable adults
## 5                People with care and support needs
## 6                           Better Care Fund
~~~

The researcher could examine what indicators constitute a domain

~~~
ftips_indicators <- indicators(ProfileID = 8, DomainID = 1000101)

## Warning: package ‘bindrcpp’ was built under R version 3.3.3
ftips_indicators %>%
        mutate(IndicatorName =
            str_trunc(as.character(IndicatorName), width = 20, “right“)) %>%
        head()

## IndicatorID     IndicatorName     DomainID     DomainName   ProfileID
## 1     1107 Total number of d… 1938132733 Better Care Fund      8
## 2     1194 ASCOF, 2A(2) - Pe… 1938132733 Better Care Fund      8
## 3     1195 ASCOF, 2C(1) - To… 1938132733 Better Care Fund      8
## 4     1196 ASCOF, 2C(2) - De… 1938132733 Better Care Fund      8
## 5    22401 2.24i - Age stand… 1938132733 Better Care Fund      8
## 6    22402 2.24ii - Age stan… 1938132733 Better Care Fund      8
##       ProfileName
## 1 Adult Social Care
## 2 Adult Social Care
## 3 Adult Social Care
## 4 Adult Social Care
## 5 Adult Social Care
## 6 Adult Social Care
~~~

The researcher may then wish to pull down the data for one or more indicators from one or more domains or profiles. However, before they can do so, they need to check what geographies are represented as the indicators data are not always available for all geographies.

~~~
ind_at <- indicator_areatypes() %>%
     left_join(area_types(), by = c(“AreaTypeID”, “AreaTypeID“)) %*>*%
     select (IndicatorID, AreaTypeID, AreaTypeName) %*>*%
     unique() head(ind_at)
##   IndicatorID AreaTypeID
## 1        108         6
## 2        108       101
## 11       108       102
## 18       113         6
## 19       113       101
## 28       113       102
##                                 AreaTypeName
## 1                     Government Office Region
## 2   Local authority districts and Unitary Authorities
## 11               Counties and Unitary Authorities
## 18                   Government Office Region
## 19 Local authority districts and Unitary Authorities
## 28               Counties and Unitary Authorities
~~~

The researcher is now in a position to read the data into the working environment in R.

~~~
ftips_data <- fingertips_data(IndicatorID = 90362, AreaTypeID = 102)
ftips_data %>%
        select(IndicatorID, IndicatorName, AreaCode, AreaName, AreaType,
      Sex, Age, Timeperiod, Value, LowerCIlimit, UpperCIlimit) %>%
        mutate(IndicatorName =
                str_trunc(as.character(IndicatorName), width = 20, “right“),
            AreaName = str_trunc(as.character(AreaName), width = 20, “right“)) %>%
        head()
## IndicatorID        IndicatorName   AreaCode   AreaName    AreaType
## 1     90362 0.1i - Healthy li… E92000001      England  Country
## 2     90362 0.1i - Healthy li… E92000001      England  Country
## 3     90362 0.1i - Healthy li… E12000001   North East region  Region
## 4     90362 0.1i - Healthy li… E12000002   North West region  Region
## 5     90362 0.1i - Healthy li… E12000003 Yorkshire and the…  Region
## 6     90362 0.1i - Healthy li… E12000004 East Midlands region Region
## Sex          Age Timeperiod   Value LowerCIlimit UpperCIlimit
## 1   Male All ages 2009 - 11 63.03181    62.88849     63.17513
## 2 Female All ages 2009 - 11 64.07049    63.92063     64.22036
## 3   Male All ages 2009 - 11 59.74471    59.22943     60.26000
## 4   Male All ages 2009 - 11 60.77587    60.41411     61.13762
## 5   Male All ages 2009 - 11 60.84513    60.39213     61.29812
## 6   Male All ages 2009 - 11 62.61274    62.08066     63.14483
~~~

### Using the package to investigate deprivation and life expectancy at birth

To expand on the functions described above, suppose a researcher wishes to examine the relationship between socio-economic deprivation and life-expectancy at birth. The fingertipsR package provides a convenient method to extract the data from the fingertips website directly into the working environment of R.

First, the researcher loads in the deprivation data at the level of the upper-tier local authorities (also described as County and Unitary Authorities). (Smith et al. 2015)

~~~
dep <- deprivation_decile(AreaTypeID = 102, Year = 2015)
head(dep)
##     AreaCode IMDscore decile
## 1 E06000001    33.178      2
## 2 E06000002    40.216      1
## 3 E06000003    28.567      3
## 4 E06000004    24.625      5
## 5 E06000005    23.639      5
## 6 E06000006    31.943      2
~~~

The researcher can then limit the previously loaded data set giving life expectancy at birth to the level of upper-tier local authority and apply the deprivation data to it.

~~~
ftips_data <- ftips_data %>%
     *# restrict to relevant geography and time*
     filter(AreaType == “County & UA” & Timeperiod == “2012 - 14“) %>%
     *# merge in deprivation data*
     left_join(., dep)
~~~

The researcher can then plot the relationship between life expectancy at birth and deprivation

~~~
p <- ggplot(ftips_data, aes(x = IMDscore, y = Value)) +
      geom_point() +
      geom_smooth(se = FALSE, method = “loess”) +
      facet_wrap(~ Sex) +
      scale_x_reverse(“IMD score”) +
      scale_y_continuous(“Life expectancy”)
p
~~~

**Figure 1.**
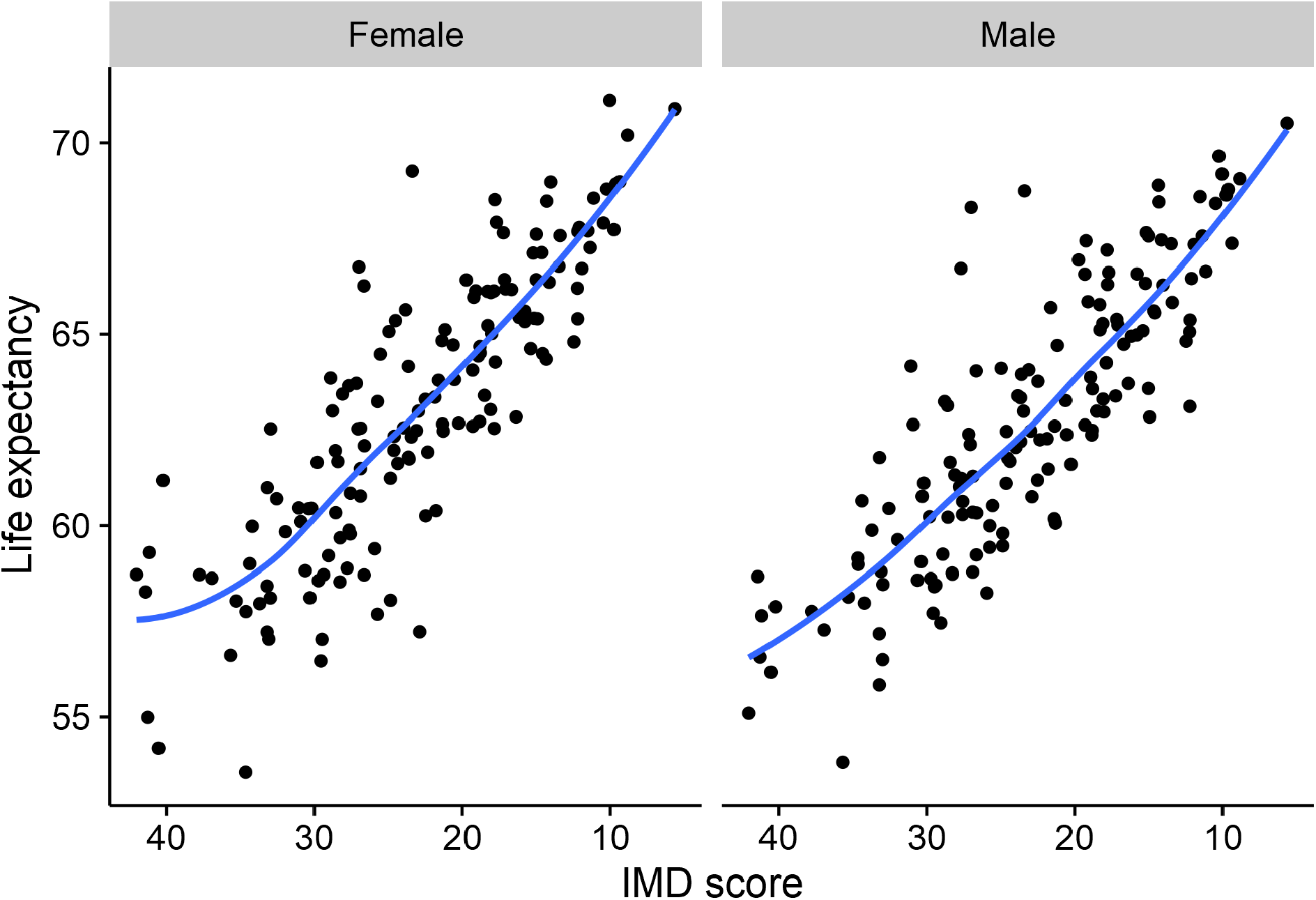
**Life expectancy at birth by Index of Multiple Deprivation score, Upper Tier Local Authority, England 2012-14**

### Extending the package

The Fingertips website, along with its API, are in continuous development. The website has new data added to it on a monthly schedule, and any changes to the API are implemented at the same time. As a result there is a lot of scope for extension to the functions that could be provided in the R package.

The package was developed openly on GitHub, an online archive where computer code is stored, and in which it is carefully version controlled through Git software. Various features of Git and GitHub facilitate scientific collaboration. (Ram 2013, Bryan (2017)) GitHub is a decentralised system - copies of code can be stored locally (i.e. on a computer hard drive) and remotely (as *forks* on the GitHub website) then synchronised at the main repository. Crucially, Git and GitHub allow any user to examine and adapt code then provide the adapted code back to the main repository for a project. Additionally, GitHub provides an issue tracker with which users of software can report problems with the software or suggest new features that might be incorporated. The inherent version control identifies all changes that were made to analytic code, who made them and at what time. This allows analytic code to be reverted to previous states, meaning that breaking changes can be easily reverted. Feedback based on the original release of the package has lead to developments to support different types of researchers, including:

- add a `find_indicator` graphical user interface, to support users less comfortable with R’s square bracket indexing
- add a function to identify areas that are significantly worse than the national average and show a significant trend towards worse values

Additional suggestions for future updates include:

- predict future indicator values based on other indicators within a profile
- detect extreme values
- flag new or recently updated indicators

Users of the fingertipsR package are encouraged to add to the issues list if they feel extensions might be beneficial for the users of this package. At the date of writing, the package itself has been downloaded from CRAN 607 times, 30 issues have been raised at GitHub and four researchers have contributed to the project repository.

## Discussion/conclusions

The work presented here illustrates a number of new and important concepts in public health research. Software for public health can be developed by researchers who are not professional software developers and can be distributed freely and easily on the web. Open source software (software where the code can be examined and adapted) allows the rapid development of new functionality, greatly expanding the uses to which it can be put and rapidly providing solutions to otherwise unmet need. The decentralised nature of the version control software Git means that scientific software and analytic code can be developed without the need for phsical co-location of collaborators - some of the authors of this paper have never physically met. Open data facilitates the sharing of information important to public health. Together, open data, open source software and distributed software development have a great deal to offer public health.

